# The Inflammatory Role of Milk Fat Globule Epidermal Growth Factor VIII in Angiotensin II Induced Arterial Remodeling

**DOI:** 10.1101/2021.03.11.434767

**Authors:** Leng Ni, Lijuan Liu, Wanqu Zhu, Richard Telljohann, Jing Zhang, Robert E. Monticone, Kimberly R. McGraw, Changwei Liu, Christopher H. Morrell, Pablo Garrido-Gil, Jose Luis Labandeira-Garcia, Edward G. Lakatta, Mingyi Wang

## Abstract

**Background:** Angiotensin II (Ang II) and milk fat globule-epidermal growth factor VIII (MFG-E8) are involved in age-associated arterial remodeling; however, the inflammatory role of MFG-E8 in Ang II associated arterial remodeling with aging remains unknown.

**Methods and Results:** In this study, 30-week-old MFG-E8 knock out (KO) and age-matched wild-type (WT) mice were infused with Ang II or saline. After infusion the with Ang II, the aortic molecular, cellular, and structural remodeling were observed in mice and compared to those infused with saline, but these effects were dependent on the expression of MFG-E8: (1) In the WT mice, Ang II infusion substantially increased intimal-medial thickness, elastic lamina degradation, collagen deposition, and the proliferation of VSMCs; in contrast, in the KO mice, these effects were significantly reduced; (2) In the WT mice, Ang II treatment significantly increased the activation and expression of MMP2, TGF-β1, and its downstream signaling molecule p-SMAD2, and collagen type I production, however, in the KO mice, these molecular effects were significantly reduced; (3) In the WT mice, Ang II treatment increased inflammatory p-NF-κB p65, MCP1, TNF-α, ICAM1, and VCAM1 molecular expression, while conversely, in the KO mice, no significant inflammatory changes were found; (4) Importantly, compared to untreated control mice with a wide range of age from 4-96 weeks, Ang II infused “younger” mice produced an “older” arterial inflammatory phenotype, which was alleviated by MFG-E8 deficiency.

**Conclusions:** MFG-E8 mediates Ang II associated arterial inflammatory remodeling. Targeting MFG-E8 is a novel molecular approach to curb adverse arterial remodeling during aging and hypertension.

**CLINICAL PERSPECTIVE:** *What Is New?:*

• Both Ang II and MFG-E8 increases are involved in proinflammatory arterial remodeling mediating the molecular, cellular and tissue events in aging and hypertension.
• MFG-E8 is essential for Ang II induced and age-associated adverse arterial remodeling via the increase of proinflammation, intimal medial thickening, elastin fragmentation, collagen deposition, and VSMC proliferation.

*What Are the Clinical Implications?:* Since MFG-E8 mediates Ang II induced proinflammation in arterial wall remodeling in aging and hypertension, targeting MFG-E8 is a potential molecular approach to curb inflammatory arterial remodeling, maintaining the health of the vascular system during aging and hypertension.

## Introduction

Angiotensin II (Ang II), a major metabolic component of the renin-angiotensin-aldosterone signaling system (RAAS), produces multiple cardiovascular inflammatory effects. ^1, 2^ Physiologically, Ang II signaling is critical in regulating blood pressure. In pathological conditions, Ang II stress signaling induces inflammation, contributing to the development of arterial aging and hypertensive vasculopathy. ^1, 3^ Our previous study showed that younger adult animals infused with Ang II exhibited an older arterial phenotype, including increased intimal vascular smooth muscle cells (VSMC) infiltration, intimal-medial thickening, matrix metalloproteinase type 2 (MMP2) activation, transforming growth factor beta 1 (TGF-β1) activation, and collagen deposition, eventually leading to elevated systolic blood pressure (SBP).

Milk fat globule-epidermal growth factor 8 (MFG-E8) is a secreted extracellular glycoprotein and that was initially discovered as a bridging molecule between apoptotic cells and macrophages for the clearance of cellular debris, also known as efferocytosis. ^5, 6^ Recent studies have also shown that MFG-E8 exerts an inflammatory role in vascular remodeling. ^7–13^ *In vivo* studies have demonstrated that MFG-E8 protein levels increase during arterial aging and injury, and in hypertensive, diabetic, and atherosclerotic arterial walls. ^7–12, 14, 15^ *In vitro* studies have demonstrated that MFG-E8 is a pivotal relay element within the Ang II/monocyte chemoattractant protein 1 (MCP1) signaling cascade that mediates VSMC inflammation, invasion, and proliferation. ^7, 8^ These *in vivo* and *in vitro* studies reveal that MFG-E8 signaling plays an important role in the initiation and progression of adverse arterial remodeling such as intimal medial thickening.

Both MFG-E8 and Ang II increase in the inflamed aging arterial wall or in cultured aging VSMCs, ^7, 8^ however, whether the underlying inflammatory role of MFG-E8 during Ang II induced adverse arterial remodeling with aging is crucial remains unknown. Therefore, we hypothesized that a chronic infusion of Ang II to younger mice, like younger rats, ^4^ would generate an “older” arterial inflammatory phenotype, which is modulated by MFG-E8 signaling.

To this end, younger (30-week-old) wild type (WT) and MFG-E8 knockout (KO) mice infused with Ang II or saline were utilized and compared to untreated control mice with a wide range of age from 4-96 weeks to investigate how MFG-E8 modulates Ang II associated arterial inflammatory remodeling at the molecular, cellular, and tissue levels with aging and hypertension.

## Materials and Methods

All data, analytic methods, and study materials are available to other researchers for purposes of reproducing the results or replicating a procedure. The material that supports the findings of this study are available from the corresponding author upon reasonable request.

### Experimental Animals

All experiments were conducted according to the protocols (445-LCS-2022) approved by the National Institute on Aging (NIA) in accordance with the National Institute of Health (NIH) Animal Care and Use Committee. MFG-E8 knock out (KO) mice that were generated, characterized, and genotyped as described previously, were obtained from Dr. Mark Udey at National Cancer Institute (NCI). ^16, 17^ Transgenic Rip1-Tag2 mice were obtained from the National Cancer Institute (NCI). MFG-E8 KO mice were generated by replacing exons 2 to 6 of the gene encoding MFG-E8 in 129SvJ embryonic cells (ES) cells with a neomycin resistant cassette. Mice were genotyped by PCR using the following primers: Rip1-Tag2, GGACAAACCACAACTAGAATGCAGTG (forward) and CAGAGCAGAATTGTGGAGTGG (reverse); Neo, GCCAGAGGCCACTTGTGTAG; and MFG-E8, CTCTCAGATTCACCTGCTCGTG and CACCGTTCAGGCACAGGCTG. 30-week-old male MFGE8 KO mice (homozygous for lack of MFG-E8) and age-matched WT mice were used in this Ang II infusion study. In addition, 4-, 8-, 20, 50-, and 96-week-old male WT and age-matched KO archival aortic sections or frozen tissue were employed for the supplemental study of age-related arterial remodeling.

### Angiotensin II Infusion and Blood Pressure Measurement

Osmotic mini-pumps (Alzet Model 2004) were implanted dorsally and subcutaneously in anaesthetized (2% isoflurane) mice (n=14-18/group) to deliver Ang II (500 ng/kg/min) or 0.9% saline (placebo) for 28 days.

SBP was measured using the pneumatic tail-cuff method (MRBP System, Life Science, Woodland Hills, California). Briefly, animals were placed in a plastic chamber maintained at 34°C and a cuff with a pneumatic pulse sensor was attached to the tail for SBP measurements. Mice were trained for one week to become accustomed to the new handling and environment. Once SBP values were consistent, eight consecutive measurements were performed for each mouse and all the values collected were averaged. Blood pressure was monitored 1 week prior to mini-pump implantation (baseline), and once per week from day 7 to 28 after pump implantation.

### Sample Collection and Preparation

#### Blood Collection

Ocular blood was collected at 14 days post mini-pump implantation. In the last week of the experiment, mice were sacrificed under sodium pentobarbital anesthesia, and blood was collected from the right atrium into a 4ml Ethylenediaminetetraacetic acid (EDTA) coated Eppendorf tube immediately. The blood samples were centrifuged for 15 minutes at 3000 rpm (1500 × g) at 4°C. The supernatant was carefully transferred into a 0.5 ml Eppendorf tube and stored at −80^°^C.

#### Tissue Collection

Tissue samples were isolated from the ascending aorta to the bifurcation of the common iliac artery. For morphological analysis, animals were perfused with 4% paraformaldehyde-phosphate buffered saline (PBS) at physiological pressure for 5 minutes; and aortic tissues were paraffin-embedded and sectioned for histological analysis. For western blotting, animals were perfused with 0.9% saline and fresh aortic tissues were snap-frozen in liquid nitrogen and stored at −80^°^ C.

### Histology, Immunostaining, and Morphometric Analyses

To quantitate aortic remodeling, IMT was measured using hematoxylin and eosin (H&E) staining; elastin breaks were counted via Elastin Verhoeff’s-Van Gieson (EVG) staining; and collagen deposition was evaluated with Masson’s trichrome staining. H&E, EVG, and Masson trichrome staining were performed using MasterTech stain kits (StatLab, McKinney, TX). Staining of aortic walls was performed as described in previous studies. ^18^ In brief, aortic paraffin sections (5 µm in thickness) were utilized for immunostaining with anti-intercellular adhesion molecule 1 (ICAM1), vascular cell adhesion molecule 1 (VCAM1), and collagen type I antibodies. Details of primary antibodies used are listed in **Table 1**. The ratios of target immunohistochemical staining positive area to the total tissue or cell area were determined via a computer-imaging program according to the instruction provided by manufacture (MetaMorph Imaging System, Molecular Devices, San Jose, CA).

**Table 1.** Primary antibodies.

| **Antibody** | **Company** | **Cat#** | **Titer for WB** | **Titer for IHC** |
| --- | --- | --- | --- | --- |
| Ang II | Sigma | A9525 |  | 1:350 |
| AT1 | Santa Cruz | sc-515884 | 1:500 | 1:100 |
| AT2 | Abcam | Ab92445 | 1:1000 | 1:100 |
| Collagen I | Santa Cruz | sc-59772 |  | 1:50 |
| GAPDH | Cell Signaling technology | 5014 | 1:1000 |  |
| ICAM1 | Santa Cruz Biotechnology | Sc-8439 |  | 1:50 |
| MCP1 | R&D Systems | AF479 | 1:200 |  |
| MFG-E8 | R&D Systems | AF2805 | 1:1000 |  |
| MMP2 | R&D Systems | AF1488 | 1:500 |  |
| MT1-MMP | Thermo Fisher Scientific | PA5-16514 | 1:500 |  |
| NF-κB p65 | Santa Cruz Biotechnology | sc-8008 | 1:500 | |
| p-NF-κB p65 | Santa Cruz | Sc-33020 | 1:500 | 1:50 |
| PCNA | Santa Cruz | sc-9857 |  | 1:50 |
| p-Smad2 | Santa Cruz Biotechnology | 3108 | 1:500 |  |
| Smad2 | Santa Cruz Biotechnology | 5339 | 1:1000 |  |
| TGF-β1 | Santa Cruz Biotechnology | sc-146 | 1:200 | |
| TNF-α | Abcam | ab9739 | 1:1000 | 1:100 |
| VCAM1 | Santa Cruz Biotechnology | Sc-13160 |  | 1:50 |

WB=western blotting; IHC=immunohistochemistry

### Western Blotting

Western blot analysis was performed as described previously. ^18^ In brief, total protein was quantified using the Pierce ^TM^ Coomassie (Bradford) protein assay kit (Hercules, CA, USA), following the manufacturer’s instructions. Ten µg of total protein was run on 4-12% NuPAGE gels (Thermo Fisher Scientific, Waltham, MA), then transferred to a polyvinylidene fluoride (PVDF) membrane and immunoblotted with the antibodies listed in **Table 1**. Western blotting bands were quantified using NIH image J (http://rsb.info.nih.gov/nih-image/) and the intensity values were normalized to loading control GAPDH or β-actin.

### PAGE Zymography

MMP-2 activity was determined via Polyacrylamide Gel Electrophoresis (PAGE) gelatin zymography (Thermo Fisher Scientific, Waltham, MA) as described previously.^19^

### Ang II Quantification

RayBio® Mouse angiotensin II ELISA Kit, a commercially available enzyme-linked immunosorbent assay (ELISA), (RayBiotech Life, Peachtree Corners, GA) was used to measure the amount of Ang II in mouse plasma.

### Statistical Analysis

All data were presented as mean ± SEM. Statistical analyses used two-way or two-way repeated measure ANOVA followed by Bonferroni post hoc test used for multiple comparisons. These statistical analyses were performed using GraphPad Prism (version 8.4.1) software. For SBP, the repeated-measures data were also analyzed using a linear mixed-effects model.^20^ A value of p≤0.05 was considered statistically significant.

## Results

### Ang II Infusion Increases Aortic MFG-E8 and Systolic Blood Pressure

Current supplemental studies (**Supplemental results and Figures I-III**) demonstrated that the vasoactive mediator Ang II, its receptors AT1/AT2, and its downstream extracellular molecules MFG-E8 and transcription factor NF-κ B played an important role in inflammatory adverse arterial remodeling, including intimal medial thickening, in WT mice. Importantly, in KO mice, these aging effects were significantly alleviated, including increases in Ang II, p-NF-κB p65, and AT1 expression and decrease in AT2 abundance (**Supplemental results and Figures I-III**).

To determine if the above effects are modulated by Ang II/MFG-E8 signaling cascade, younger WT and KO mice (30-week-old) infused Ang II or saline were utilized. Ang II infusion markedly increased the levels of circulating Ang II in both WT and KO mice on day 14 and day 28, compared to saline infused animals. Unexpectedly, a significant higher of Ang II levels was observed in KO vs. WT (**Figure 1A**). Importantly, Ang II infusion dramatically increased the expression of aortic MFG-E8 in WT but that aortic MFG-E8 protein was not detected in the KO mice infused with either Ang II or saline (**Figure 1B & C**). Notably, the AT1/AT2 receptor relays the signaling derived from Ang II in the arterial wall. Western blotting indicated that aortic MFG-E8 protein levels were markedly decreased in the AT1 KO, but the levels were not altered in AT2 KO mice **(Supplemental Figure IV).**

**Figure 1.**
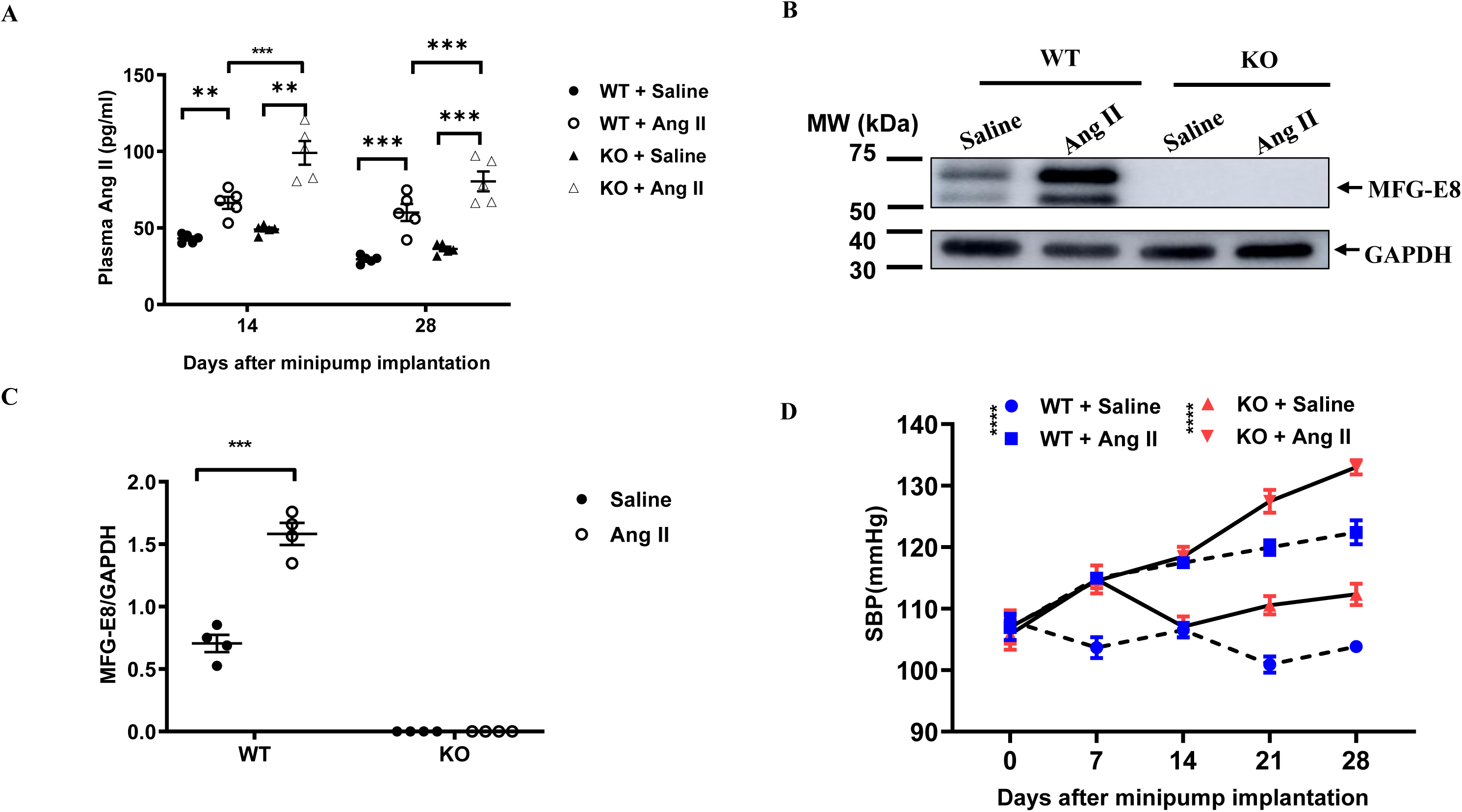
Ang II infusion increases circulating levels of Ang II and aortic MFG-E8 protein and systolic blood pressure. **A.** Graph shows plasma Ang II concentrations (p<0.001 for overall treatment effect by repeated two-way ANOVA with the factors of treatment and genotype). Graph shows mean ± SEM with individual animal data from KO and WT mice with Ang II or saline infusion over time. **=p<0.01 and ***=p<0.001 by Bonferroni post-hoc tests following two-way ANOVA. **B.** Representative Western blots of aortic MFG-E8. **C**. Quantitative data of Western blot of aortic MFG-E8, graph shows MFG-E8 protein abundance as normalized by GAPDH (p<0.001 for main Ang II infusion effect, p<0.001 for main genotype effect, p<0.001 for treatment x genotype, by two-way ANOVA). Graph shows mean ± SEM with individual data for KO and WT mice with Ang II or saline infusion. ***=p<0.001 by Bonferroni post-hoc tests following two-way ANOVA with the factors of treatment and genotype. **D.** Dynamic changes in mouse systolic blood pressure (SBP) measured by tail-cuff method; plot shows mean ± SEM for KO and WT mice (n=5/group) with Ang II or saline infusion over time (p<0.001 for main treatment effect, p<0.001 for main genotype effect, p<0.001 for Ang II infusion x genotype, by repeated two-way ANOVA and a linear mixed-effects model). ****=p <0.0001 SBP in KO + Ang II vs KO + saline and WT + Ang II vs WT + saline group.

Increased SBP induced by Ang II infusion was observed in all mice. The repeated, two-way ANOVA analysis showed that SBP had a statistically significant of treatment x genotype interaction (p<0.0001) (**Figure 1D).** At baseline (time=day 0) there was no significant difference in SBP among any of the groups. However, after Ang II infusion, both WT and KO, showed significantly increased SBP levels compared to saline infused mice. Unexpectedly, in KO, the SBP of Ang II infused mice was significantly elevated compared to WT after the infusion (p<0.01); and similarly, in KO, the SBP of saline infused was also significantly elevated compared to WT after the infusion (p<0.01) (**Figure 1D)**. However, the percentage change of the SBP after Ang II infusion was not significantly altered in KO vs WT (**Supplemental figure V**), suggesting that Ang II infusion produced similar hypertensive effect.

In addition, the baseline blood test results from 10-week-old male WT and KO mice showed that the absence of MFG-E8 significantly decreased plasma cholesterol and triglyceride levels but did not alter plasma glucose levels (**Supplemental Figure VI**), suggesting that MFG-E8 is involved in lipid metabolism, and may also influence Ang II metabolism, blood pressure, and the vascular remodeling.

### Ang II Infusion Activates Aortic MMP2

Increases in circulating Ang II and blood pressure enhance the activity of MMP2. ^4^ PAGE zymography and Western blotting analysis were performed to determine whether the interactive effects of treatment and genotype on the expression and activity of the gelatinase MMP2. Zymography demonstrated that the levels of activated MMP2 within the aortic walls of mice after Ang II infusion were dependent upon the presence of MFG-E8 (**Figure 2A)**: in WT mice, Ang II infusion markedly increased the levels of activated aortic MMP2 compared to saline; in KO mice, activated MMP2 in Ang II infused animals was significantly higher than saline, but was significantly less than that of the WT Ang II treated (**Figure 2A)**. Western blot analysis showed that the abundance of aortic MMP2 protein induced by Ang II was dependent upon the presence of MFG-E8 (**Figure 2B):** in WT mice, Ang II markedly increased the levels of MMP2 protein in aortic walls when compared to saline **(Figure 2B middle panel)**; in KO mice, the MMP2 protein levels were significantly increased in Ang II versus saline treated mice but was significantly less than that of the WT Ang II treated mice **(Figure 2B middle panel)**. Similarly, Ang II increased and induced the levels of member type I of MMP activator (MT1MMP), an activator of MMP2, was dependent on the presence of MFG-E8 (**Figure 2B right panel)**; in WT mice, Ang II significantly increased the levels of MT1MMP protein in the aortic wall compared to saline; however, in KO animals, there was no significant increase of MT1MMP protein levels (**Figure 2B right panel)**. MFG-E8 mediates the increased activation and expression of MMP2 induced by Ang II in the aortic walls.

**Figure 2.**
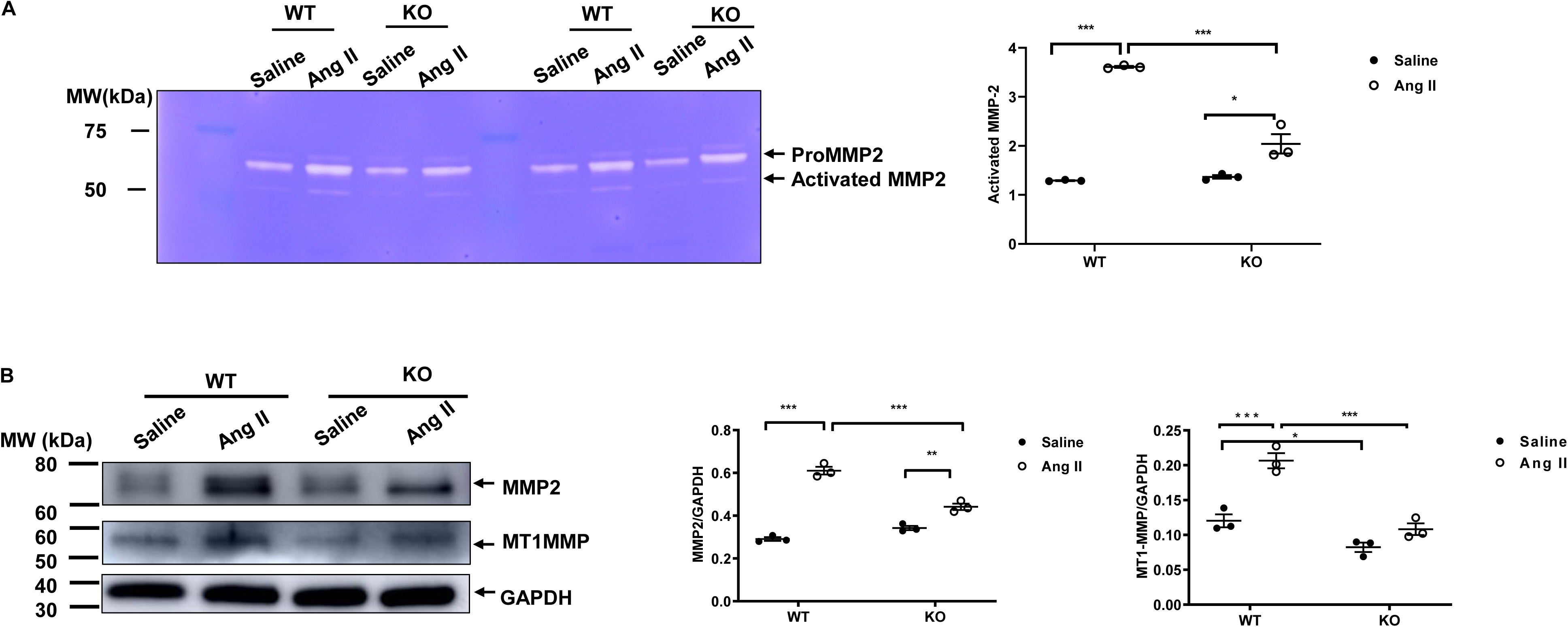
Ang II infusion enhances the activation of aortic MMP2 in WT mice. **A.** Representative zymograms of aortic gelatinases (**left panel**). Quantitative graph (**right panel**) shows activated MMP2 protein abundance (p<0.001 for main treatment effect, p<0.001 for main genotype effect, p<0.001 for treatment x genotype, by two-way ANOVA). Graph shows mean ± SEM with individual data for KO and WT mice with Ang II or saline infusion. *p<0.05 and ***=p<0.001 by Bonferroni post-hoc tests following two-way ANOVA with the factors of treatment and genotype. **B.** Representative Western blots of aortic MMP2, MT1MMP, and GAPDH (loading control). Graph shows quantitative data of (middle panel) aortic MMP2 protein abundance (p<0.001 for main treatment effect, p<0.01 for main genotype effect, p<0.01 for treatment x genotype, by two-way ANOVA). Graph shows quantitative data of (right panel) aortic MT1MMP2 protein abundance (p<0.001 for main treatment effect, p<0.01 for main genotype effect, p<0.01 for treatment x genotype, by two-way ANOVA). Graph shows mean ± SEM with individual data for KO and WT mice with Ang II or saline infusion. *=p<0.05, **=p<0.01, and ***=p<0.001 by Bonferroni post-hoc tests following two-way ANOVA with the factors of treatment and genotype.

### Ang II Infusion Increases Aortic TGF-β1 Fibrotic Signaling

Ang II signaling and its subsequent blood pressure elevation, as well as increased MMP2 activation, all promote the activation of transforming growth factor-beta 1 (TGF-β1) and its fibrotic effect within arterial walls *in vivo* and in cultured VSMCs. ^4, 21, 22^ Western blotting analysis demonstrated that the Ang II-induction of activated arterial TGF-β1 was dependent upon the presence of MFG-E8 (**Figure 3A & 3B)**: in WT mice, Ang II infusion markedly increased the levels of activated TGF-β1 protein in the aortic walls; while in KO mice there was a smaller yet significant increase when compared to saline (**Figure 3A & 3B)**. Notably, in Ang II infused mice, the levels of TGF-β1 protein in the WT was significantly higher than those found in KO (**Figure 3A & 3B)**. Similarly, Western blot analysis demonstrated that Ang II infusion increased SMAD2 phosphorylation (p-SMAD2), a TGF-β1 downstream signaling molecule, and was also dependent upon the presence of MFG-E8 (**Figure 3A & 3B).** In WT mice, Ang II infusion markedly increased the levels of activated p-SMAD2 protein in the aortic walls compared to saline; in contrast, in KO mice there was no significant increase of these proteins with Ang II infusion (**Figure 3A & 3B)**. Expectedly, immunohistostaining and Western blotting analyses indicated that the Ang II induction of the arterial collagen I levels, a TGF-β1/p-SMAD2 downstream signaling product, were dependent upon MFG-E8 (**Figure 3C & 3D).** In WT mice, Ang II infusion markedly increased the levels of collagen type I protein in the aortic walls compared to saline infusion; in contrast, in KO mice there was no significant increase of collagen type I when treated with Ang II **(Figure 3C & 3D)**. Clearly, MFG-E8 plays an important role in the Ang II induced activation of TGF-β1, its downstream signaling molecule, p-Smad2, and collagen I in the aortic walls.

**Figure 3.**
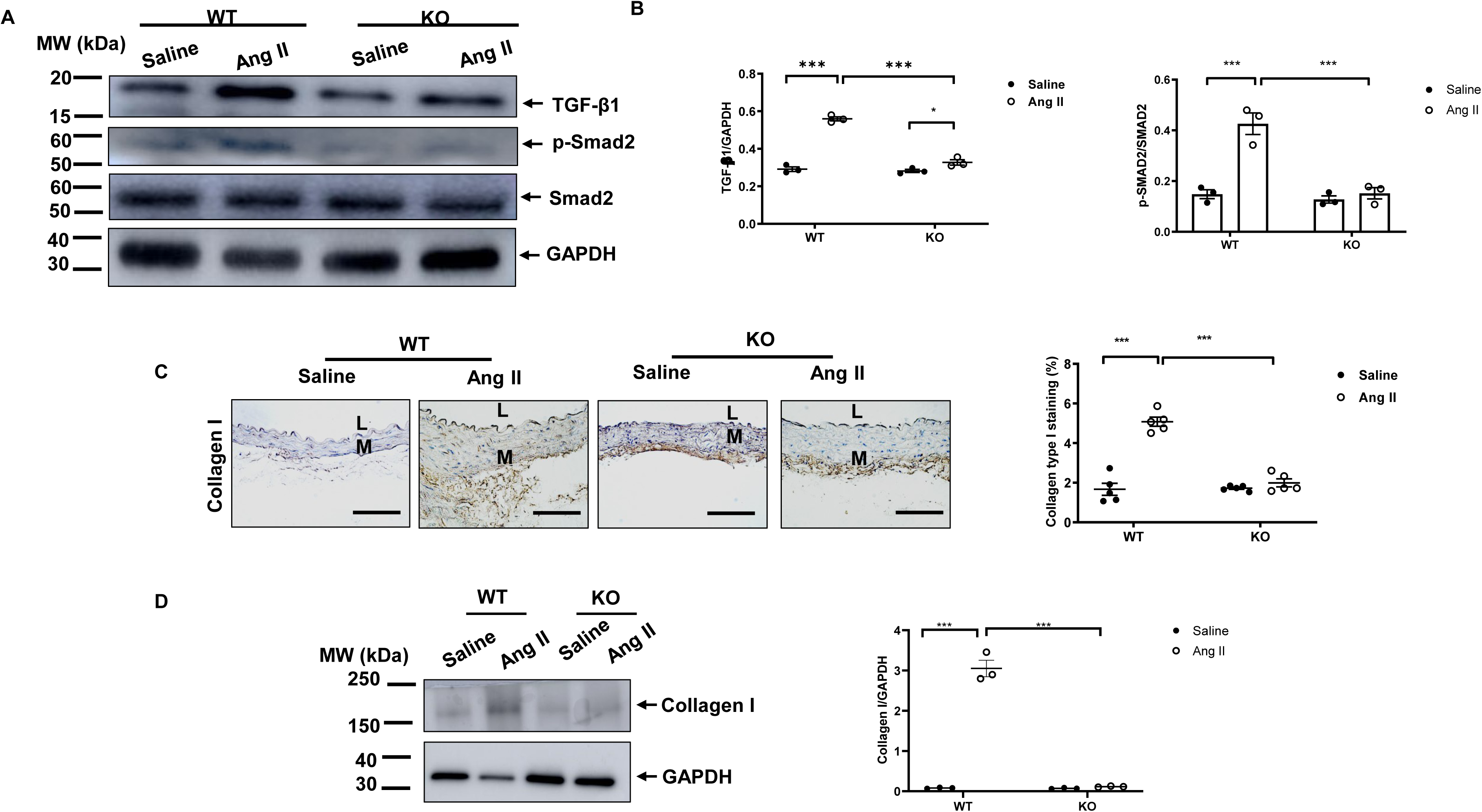
Ang II infusion increases TGF-β1 activity and signaling within the aortic walls. **A.** Representative Western blots of aortic TGF-β1, p-SMAD2, and SMAD2. **B.** Graph shows quantitative data of (middle) aortic activated TGF-β1 protein abundance (p<0.001 for main treatment effect, p<0.001 for main genotype effect, p<0.001 for treatment x genotype, by two-way ANOVA). Graph shows quantitative data of (right) aortic p-SMAD2/SMAD2 ratio (p<0.001 for main treatment effect, p<0.001 for main genotype effect, p<0.01 for treatment x genotype, by two-way ANOVA). Graph shows mean ± SEM with individual data for KO and WT mice with Ang II or saline infusion. *=p<0.05 and ***=p<0.001 by Bonferroni post-hoc tests following two-way ANOVA with the factors of treatment and genotype. **C**. Photomicrographs of immunostaining collagen type I (Collagen I) and morphometric analysis (right panel) shows relative collagen I immunostaining area (%) (p<0.001 for main treatment effect, p<0.001 for main genotype effect, p<0.001 for treatment x genotype, by two-way ANOVA). ***=p<0.001 by Bonferroni post-hoc tests following two-way ANOVA with the factors of treatment and genotype. **D.** Representative Western blots of aortic Collagen I (low left panel). Graph shows quantitative data of (low right panel) aortic Collagen I (p<0.01 for main treatment effect, p>0.05 for main genotype effect, p<0.0001 for treatment x genotype, by two-way ANOVA). Graph shows mean ± SEM with individual data points for KO and WT mice with Ang II or saline infusion. ***=p<0.001 by Bonferroni post-hoc tests following two-way ANOVA with the factors of treatment and genotype. L=lumen; M=media. Scale bar= 100µm.

### Ang II Infusion Activates NF-**κ**B Signaling and Its Downstream Proinflammatory Effectors

NF-κB is a prominent proinflammatory transcription factor in inflamed arterial walls. ^23, 24^ NF-κB p65, an element of NF-κB activation, facilitates the upregulation of its downstream inflammatory effectors MCP1, tissue necrosis factor-alpha 1 (TNF-α), ICAM1, and VCAM1 which are directly involved in the process of arterial inflammation.^24–26^

Western blot analyses showed that the abundance of arterial p-NF-κB p65 in mice infused with Ang II was dependent on the presence of MFG-E8 (**Figure 4**): p-NF-κB p65 protein abundance was markedly upregulated in WT mice infused with Ang II versus saline, whereas the protein levels remained at a similar level in both groups of KO animals. Western blotting analysis indicated that the abundance of aortic MCP1 protein induced by Ang II in mice was also dependent upon the presence of MFG-E8 (**Figure 4A)**; WT mice infused with Ang II showed a significant increase in the levels of activated MCP1 protein in the aortic wall when compared to saline; conversely, KO mice infused with Ang II exhibited slightly decreased levels of MCP1 protein expression when compared to saline (**Figure 4A).** Notably, when mice were treated with Ang II, MCP1 protein levels were significantly lower in KO versus WT (**Figure 4A**). Western blotting and immunostaining analyses also revealed that Ang II infusion altered TNF-α levels in mice and was dependent upon MFG-E8 (**Figure 4A & B)**; the levels of TNF-α protein protein in WT mice was significantly increased in the aortic wall when infused with Ang II compared to saline; however, the protein levels in KO mice were not significantly altered in either treatment group (**Figure 4A & B).** In addition, immunohistostaining and morphometric analysis demonstrated that the increased levels of arterial adhesive molecules ICAM1 and VCAM1 protein in mice treated with Ang II were also dependent upon MFG-E8 **(Figure 4C & D).** In WT mice Ang II infusion markedly increased the protein levels of both ICAM1 and VCAM1 in the aortic wall compared to saline, however, in KO mice Ang II infusion did not significantly alter either ICAM1 or VCAM1 protein abundance versus saline (**Figure 4C & D).** MFG-E8 mediates Ang II induced increases in both MCP-1 and TNF-α, and VCAM1 protein expression in aortic walls.

**Figure 4.**
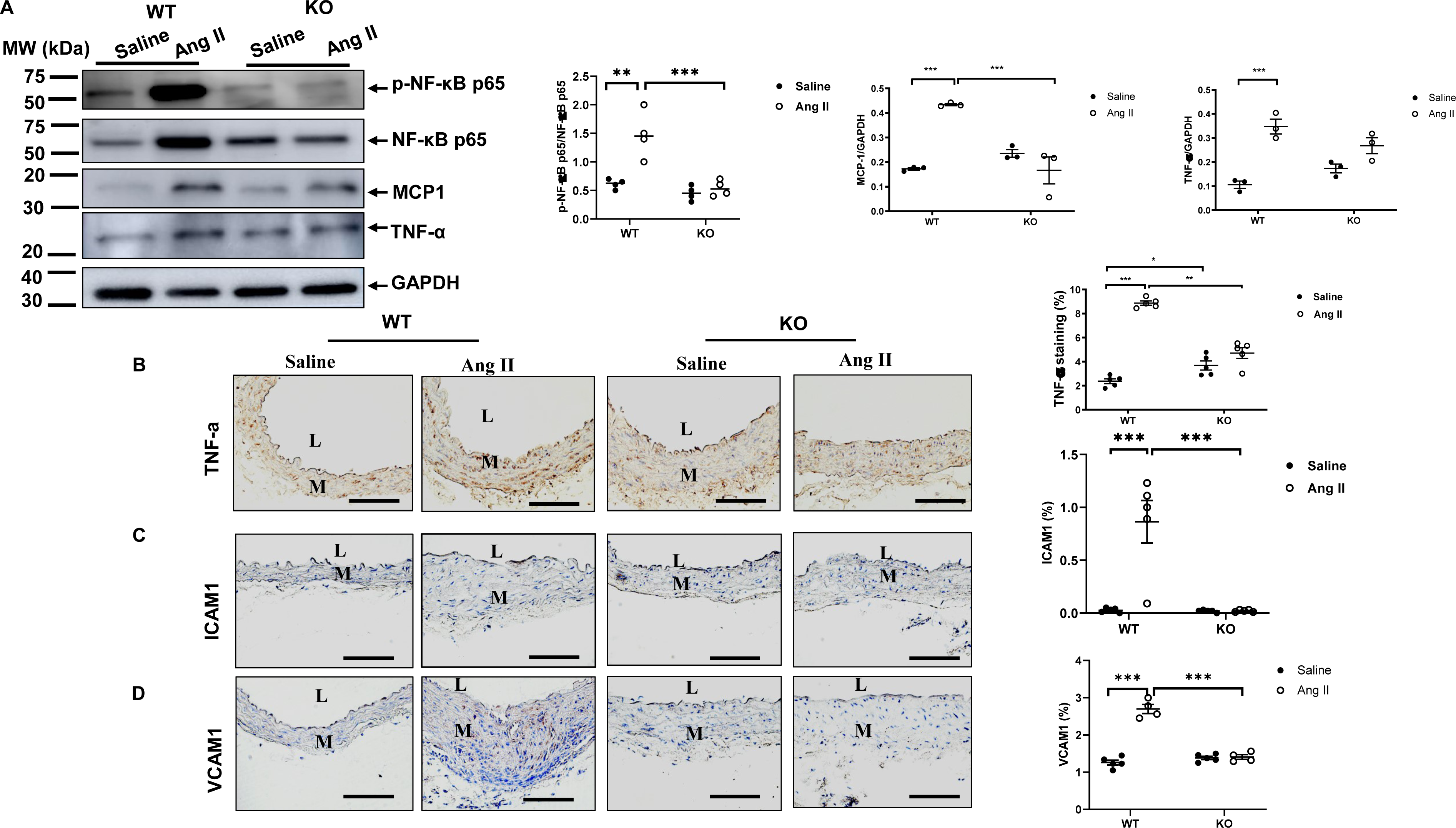
**Ang II infusion enhances aortic transcription factor NF-**κ**B activation and its downstream molecules. A.** Representative Western blots of aortic p-NF-κB p65, p-NF-κB p65, MCP-1, TNF-α, and GAPDH (left panels). Graph (second left panel) shows quantitative data of p-NF-κB p65 protein abundance (p<0.001 for main treatment effect, p<0.01 for main genotype effect, p<0.001 for treatment x genotype, by two-way ANOVA). Graph shows mean ± SEM and individual data points for KO and WT mice with Ang II or saline infusion. **=p<0.01 and ***=p<0.001 by Bonferroni post-hoc tests following two-way ANOVA with the factors of treatment and genotype. Graph (second right panel) shows quantitative data of MCP1 protein abundance (p<0.01 for main treatment effect, p<0.05 for main genotype effect, p<0.001 for treatment x genotype, by two-way ANOVA). Graph shows mean ± SEM for KO and WT mice with Ang II or saline infusion (n=5/group). ***=p<0.001 by Bonferroni post-hoc tests following two-way ANOVA with the factors of treatment and genotype. Graph (far right panel) shows quantitative data of TNF-α protein abundance (p<0.001 for main treatment effect, p>0.05 for main genotype effect, p<0.05 for treatment x genotype, by two-way ANOVA). Bar graph show mean ± SEM for KO and WT mice with Ang II or saline infusion (n=5/group). ***=p<0.001 by Bonferroni post-hoc tests following two-way ANOVA with the factors of treatment and genotype. **B**. Photomicrographs of immunostaining of TNF-α. Graph shows quantitative data of (right panel) aortic TNF-α immunostaining area (%) (p<0.01 for main treatment effect, p<0.05 for main genotype effect, p<0.01 for treatment x genotype, by two-way ANOVA). Graph shows mean ± SEM for KO and WT mice with Ang II or saline infusion (n=5/group). *=p<0.05. **=p<0.01, and ***=p<0.001 by Bonferroni post-hoc tests following two-way ANOVA with the factors of treatment and genotype. **C.** Photomicrographs of immunostaining ICAM1 and morphometric analysis (right panel) shows relative ICAM1 immunostaining area (%) (p<0.001 for main treatment effect, p<0.001 for main genotype effect, p<0.001 for treatment x genotype, by two-way ANOVA). ***=p<0.001 by Bonferroni post-hoc tests following two-way ANOVA with the factors of treatment and genotype. **D.** Photomicrographs of immunostaining VCAM1 and graph (right panel) shows relative VCAM1 immunostaining area (%) (p<0.001 for main treatment effect, p<0.001 for main genotype effect, p<0.001 for treatment x genotype, by two-way ANOVA). ***=p<0.001 by Bonferroni post-hoc tests following two-way ANOVA with the factors of treatment and genotype. L=lumen; M=media. Scale bar= 100µm.

### Ang II Infusion Induces Arterial Cellular and Matrix Remodeling

Immunohistochemistry of mouse aortic walls was performed after Ang II or saline infusion. The morphometrical analysis indicated that adverse arterial remodeling was dependent upon the presence of MFG-E8, which may be related to inflammatory molecular remodeling and changes in SBP (**Figure 1-4**): in WT mice a marked increase in IMT was observed in the aortic walls of mice infused with Ang II compared to saline; however, in KO mice, no significant difference in IMT was observed (**Figure 5A)**. Unexpectedly, after saline infusion, a significant increase in IMT was observed in KO versus WT mice (p<0.01). In WT mice treated with Ang II, EVG staining indicated a significant increase in the number of the elastin fiber breaks versus saline, while this effect was substantially reduced in KO mice (**Figure 5B**). In WT mice infused with Ang II, Masson’s trichrome staining showed that the fraction of intimal-medial extracellular matrix (ECM), mainly collagen, was increased versus saline, while this effect was not observed in KO mice (**Figure 5C**). VSMC proliferation is a key cellular event of arterial thickening in hypertension induced by Ang II. ^27^ In WT mice infused with Ang II, the percentage of proliferating cell nuclear antigen (PCNA) positive VSMCs, an index of cellular division, was markedly elevated when compared to saline; however, no difference in PCNA was seen in KO animals (**Figure 5D)**. Thus, MFG-E8 plays an important role in Ang II induced intimal medial thickening, elastic lamina degradation, collagen deposition, and proliferation of VSMCs; and notably MFG-E8 deficiency alleviates or abolish theses effects in aortic walls.

**Figure 5.**
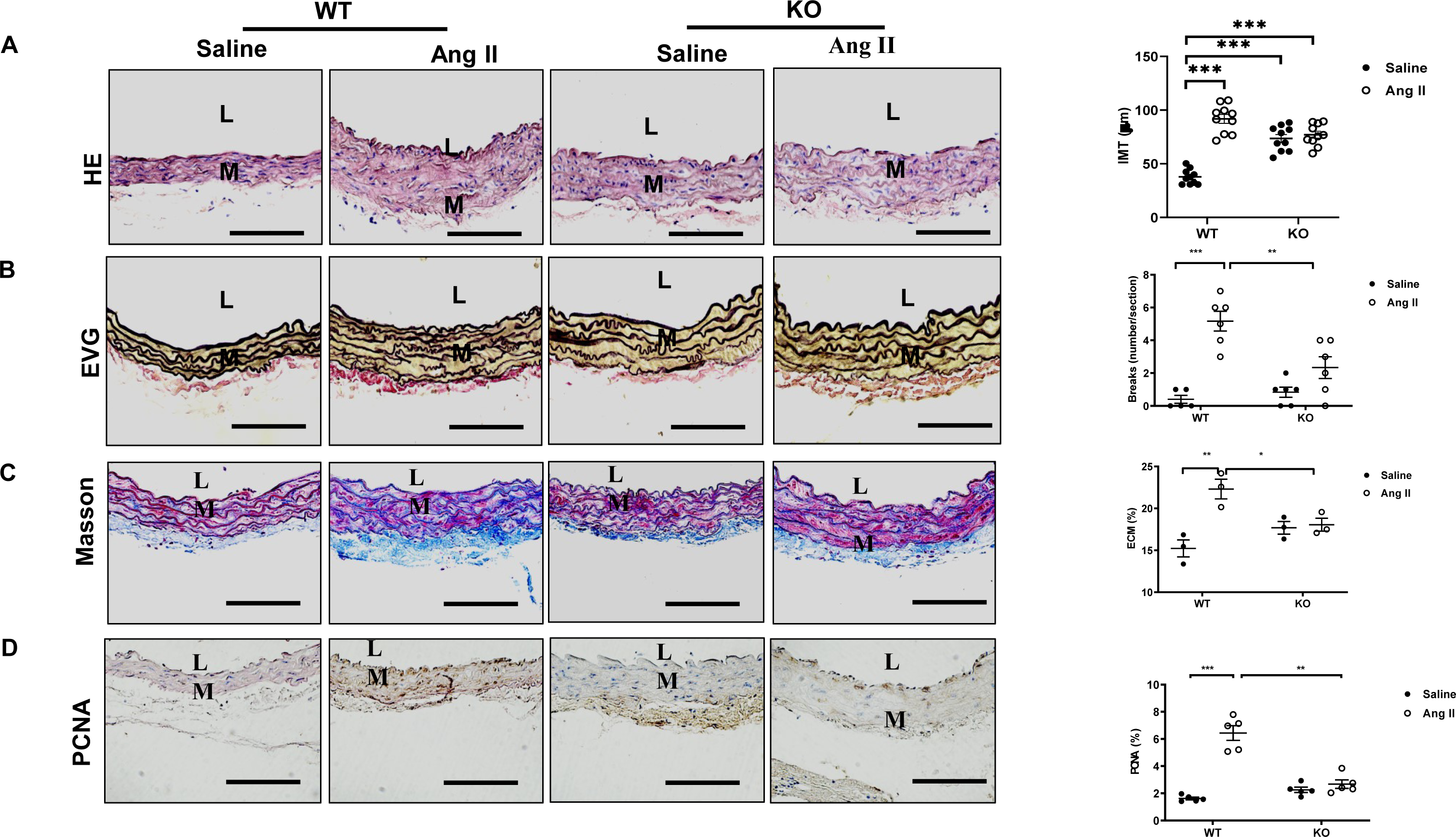
Ang II infusion increases aortic remodeling. **A.** Photomicrographs of hematoxylin-eosin (HE) staining of aortic walls. Graph shows (far right panel) the ratio of aortic intimal medial area to luminal area (p<0.001 for main treatment effect, p<0.001 for main genotype effect, p<0.001 for treatment x genotype, by two-way ANOVA). **B.** Photomicrographs of Elastin Verhoeff’s-Van Gieson (EVG staining) of aortic walls. Graph shows (far right panel) elastin breaks (p<0.05 for main treatment effect, p<0.001 for main genotype effect, p<0.01 for treatment x genotype, by two-way ANOVA). **C.** Photomicrograph of Masson’s trichrome staining of aortic walls. Graph shows intimal medial extracellular matrix (ECM) (blue color) (p>0.05 for main treatment effect, p<0.01 for main genotype effect, p<0.01 for treatment x genotype, by two-way ANOVA). **D.** Photomicrographs of proliferation cellular nuclear antigen (PCNA) immunostaining of aortic walls. Graph shows (far right panel) relative PCNA stained area (p>0.05 for main treatment effect, p<0.01 for main genotype effect, p<0.01 for treatment x genotype, by two-way ANOVA). Graph show mean ± SEM with individual animal data points for KO and WT mice with Ang II or saline infusion (n=5/group). *=p<0.05, **=p<0.01 and ***=p<0.001 by Bonferroni post-hoc tests. L=lumen; M=media. Scale bar= 100µm.

## Discussion

Ang II and MFG-E8 play an important role in inflammatory arterial remodeling in rats, nonhuman primates and humans with advancing age. ^7, 8, 12^ Current findings demonstrate that MFG-E8 mediates Ang II induced inflammatory arterial remodeling in mice (**Figure 6**). The absence of MFG-E8 alleviates Ang II induced inflammatory remodeling at the molecular, cellular, and tissue levels (**Figure 6**).

**Figure 6.**
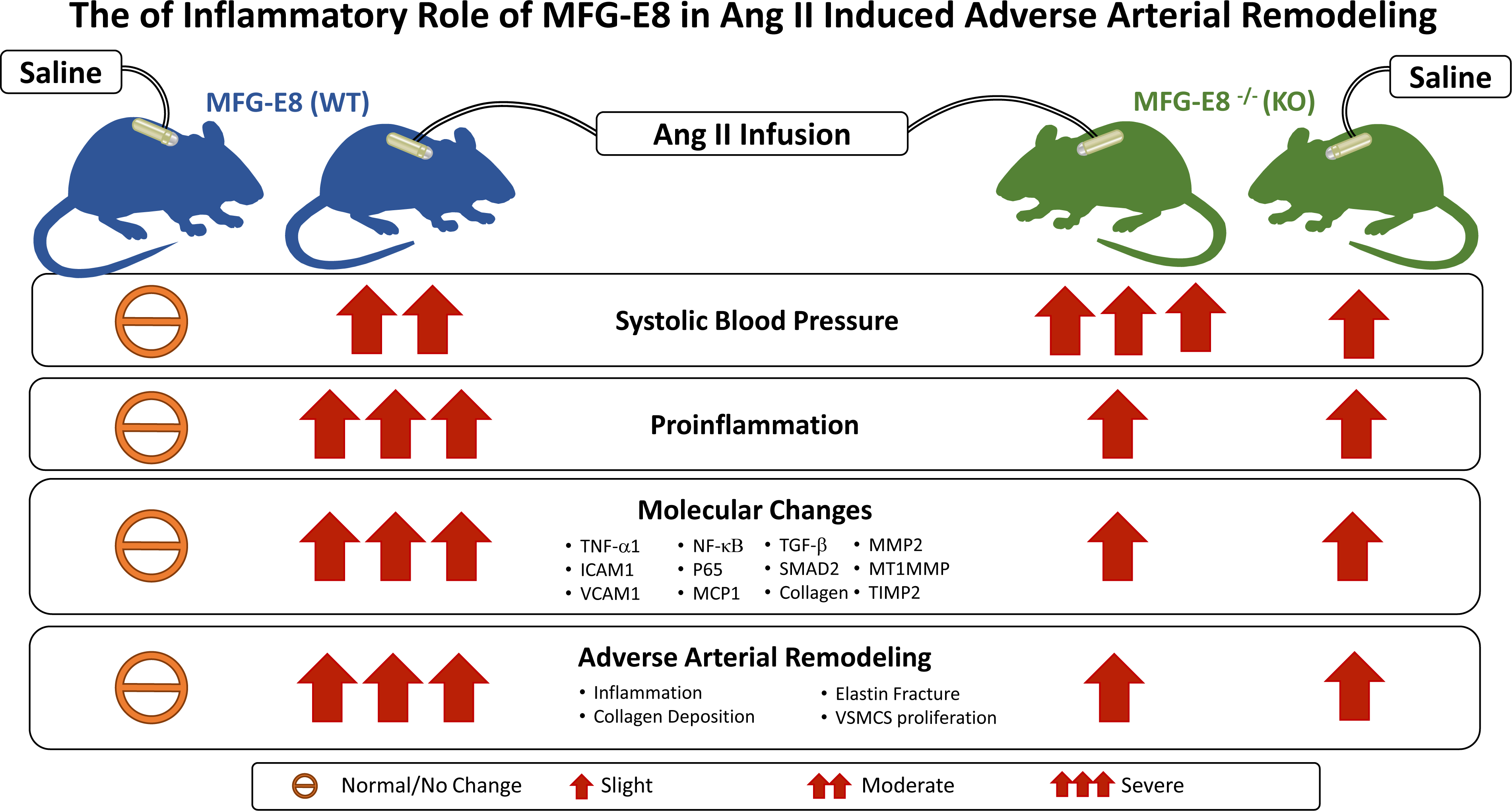
**The schematic illustration of the inflammatory role of MFG-E8 in Ang II induced arterial inflammatory remodeling.**

Ang II signaling is increased during adverse arterial remodeling with aging, which is dependent on MFG-E8 abundance. The AT1 receptor promotes Ang II inflammatory events while AT2 receptor blocks Ang II induced inflammation. ^28–30^ Current findings demonstrate that aortic Ang II and AT1 protein are increased while AT2 abundance are decreased, but these effects are alleviated by the absence of MFG-E8 in aged mice. Ang II infusion markedly increases AT1 receptor abundance while decreases AT2 expression; MFG-E8 deficiency significantly alleviates these effects. Importantly, MFG-E8 protein levels are markedly decreased in AT1 KO mice but are not significantly changed in the AT2 KO compared to WT mice. Remarkably, the absence of MFG-E8 alleviates the increase in the master inflammatory transcription factor NF-κB and intimal medial thickening in aged mice. Taken together, these findings suggest that MFG-E8 deficiency alleviates Ang II/AT1 signaling cascade-associated inflammatory arterial remodeling with advanced age. However, the detailed interactions between MFG-E8 and the receptors AT1, and AT2 during Ang II induced arterial remodeling needs to be further investigated and confirmed by pharmacologic and gene silencing approaches.

MFG-E8 are involved in age-associated change in blood pressure in mice. Aortic MFG-E8 is increased in renal hypertensive rats. ^31^ Ang II infusion increases MFG-E8 in the aortae of mice. Increased Ang II is colocalized with elevated MFG-E8 within old human aortic walls^8^. These findings suggest that MFG-E8 may closely associated with age-associated change in SBP.

Indeed, our recent findings indicate that MFG-E8 mediates the change in SBP with advancing age: there is a tendency of an increase in SBP between age 20 and 80 weeks while a decrease in SBP at age of 96 weeks in KO versus WT animals. ^32^ Current study indicates that SBP is increased over a 28-day time course in 30-week-old KO versus WT mice after saline infusion; similarly, SBP is also increased over a 28-day time course in 30-week-old KO versus WT animals after Ang II infusion. However, the percentage change of SBP is not altered in KO versus WT infused Ang II. These findings suggest that 1) MFG-E8 nonlinearly effects on age-associated change in SBP: a decrease of SBP in young adult while an increase of SBP in old mice; and 2) MFG-E8 does not have net effect but has overall effect on Ang II infusion associated SBP increase in mice **(Figure 6)**. It is well-known that MFG-E8 is a bridging molecule for the clearance of cellular debris by macrophages, and the absence of MFG-E8 impairs the clearance of cellular debris, which potentially accumulate and may subsequently cause blood pressure in young adult mice. ^11, 33^ In addition, aging increases MFG-E8 and increases its small, cleaved fragment medin, which is the most common amylogenic protein in aged arterial walls and may potentially induces inflammation and blood pressure increase in old mice. 2, 11, 34, 35

Ang II infusion induces the activation of aortic MMP2 and TGF-ß1, which promotes arterial elastic fiber degeneration and fibrosis and is dependent on MFG-E8. The arterial MMP2 activation is markedly increased in Ang II infused WT vs. KO mice, due to an increase of the MT1MMP, an activator of MMP2. We previously observed that treating VSMCs with recombinant human MFG-E8 (rhMFG-E8) activates MMP2, ^36, 37^ suggesting that MFG-E8 is a modulator of Ang II associated MMP2 activation. Notably, active MMP2 has a high capacity to degrade aortic elastin laminae and subsequently releases activated TGF-ß1. ^4, 21, 22, 38^ Ang II infusion induces greater arterial TGF-ß1 activation in the aortic walls of WT than in the MFG-E8 deficient KO mice, revealing that MFG-E8 signaling plays an important role in Ang II induced TGF-ß1 activation. In addition, MMP2, a potent activator of latent TGF binding protein (LTBP), facilitates the conversion from the latent to the active form of TGF-ß1^21^. We have previously observed that treating VSMCs with rhMFG-E8 increases the fibrogenic TGF-ß1/SMAD2/collagen type I signaling cascade in VSMCs .^36^ Thus, MFG-E8 mediates Ang II-induced MMP2 activation/TGF-β1 fibrogenic signaling during adverse arterial remodeling. ^21, 39^

Ang II infusion increased NF-κB signaling which elevates inflammation, which is dependent upon MFG-E8. NF-κB p65 is a core element of NF-κB activation during inflammatory arterial wall remodeling. ^40^ The current findings show that Ang II treatment markedly increases p-NF-κB p65 in WT but not in KO, suggesting that MFG-E8 is an important intermediary of Ang II induced NF-κB signaling. Several studies indicate that NF-κB activation facilitates the production of MCP1, TNF-α ICAM1, and VCAM1 during the processes of arterial inflammation and remodeling. ^14, 24–26, 41–45^ MFG-E8 increases MCP1 activation in VSMCs. ^8^ MCP1 and TNF-α are prominent inflammatory factors in arterial walls. ^46^ The upregulation of MCP1 and TNF-α are higher in WT mice than in KO mice infused with Ang II further suggesting that MFG-E8 is involved in Ang II induced arterial inflammation. Ang II also induced an increase in the levels of arterial ICAM1 and VCAM1 in mice, which is dependent on MFG-E8. Thus, Ang II increases NF-κB and its downstream molecules MCP1, TNF-α, ICAM1 and VCAM1, which is mediated by MFG-E8, facilitating arterial inflammation.

Ang II signaling mediated increase in IMT is modulated by MFGE8. The AT1 receptor and IMT are decreased in KO versus WT mice at age of 96 weeks. The abundance of AT1 receptor and IMT are markedly increased in 30-week-old Ang II infused versus saline-infused WT animals but are not changed in Ang II infused versus saline infused KO animals. These findings suggest that Ang II signaling mediated increased in IMT is modulated by MFG-E8 in Ang II-infused 30-week-old and untreated 96-week-old animals. In addition, it is known that both increased SBP and enhanced Ang II signaling play an important role in intimal medial thickening in rats.^4^ The increase in SBP over a 28-day time course likely contributes to the increase in IMT in KO versus WT at age of 30-week-old after saline infusion. These findings suggest that the increase of SBP over a 28-day time course in 30-week-old KO vs WT after saline infusion while the deceases in SBP and Ang II signaling AT1 receptor in untreated 96-week-old KO versus WT may contribute to the discrepancy of the findings in **Figure 5A** versus **supplement Figure 1 B** in terms of IMT of KO versus WT. However, the detailed underlying causes of this discrepancy need to be further explored in the future.

Importantly, Ang II infusion of WT mice promotes adverse arterial remodeling, including inflammation, collagen deposition, elastic fiber degradation, proliferation of VSMCs, which is mediated by MFG-E8. MFG-E8 deficiency substantially alleviates Ang II induced adverse arterial remodeling including inflammation, collagen deposition and elastic fiber breakdown in KO vs. WT. Previous and current studies indicate that increases in inflammation, MMP2 activation, TGF-β fibrotic signaling, proliferation of VSMCs are the molecular and cellular mechanisms underlying Ang II induced adverse arterial remodeling. ^2, 7–10, 12, 13, 19, 22, 46, 47^.

There is an apparent contradiction of our current findings which shows MFG-E8 in an inflammatory role compared to prior research which suggests that MFG-E8 exerts an anti-inflammatory or anti-fibrotic role. ^48–50^ For example, MFG-E8 attenuates Ang-II induces atrial fibrosis and vulnerability to atrial fibrillation through the inhibition of the TGF-β1/Smad2/3 pathway. ^48–50^ It also allevaites ventricle fibrosis via the attenuation of endothelial-mesenchymal transition through the Smad2/3-snail signalling pathway and restoring circulating MFG-E8 levels attenuates cardiac hypertrophy through the inhibition of the Akt (protein kinase B, PKB) pathway.^48–50^ In addition, MFG-E8 has biologic effects on adaptive immunity, including anti-inflammatory effects to postischemic cerebral injury, and to post lipopolysaccharide treatment.^51, 52^ Alternatively, an excess of MFG-E8 with age also increases its small, cleaved fragment medin, which potentially induces proinflammatory endothelial activation. ^34, 35^ It is well-known that medin is the most common amylogenic protein found in aged arterial walls. ^2, 11^ In the present study, Ang II induces an older phenotype which could create a condition where excess MFG-E8 is enzymatically cleaved into medin which could impair the structure and function of arterial walls. Medin associated inflammation and fibrosis in the arterial wall likely contribute to the discrepancies of MFG-E8 effects between aortic walls and the myocardium and needs to be further studied. ^34, 35, 51^

In addition, MFG-E8 is involved in metabolism, which may affect arterial inflammation. MFG-E8 deficiency significantly decreases cholesterol and triglyceride levels and has been shown to impede the development of obesity by inhibiting the uptake of dietary fats and serum fatty acids in mice.^53^ Notably, circulating levels of Ang II post-infusion are significantly higher in KO than WT animals also suggesting that MFG-E8 may be involved in the metabolism of Ang II. The underlying mechanism and inflammatory consequence of MFG-E8 in lipid profile and circulating Ang II levels are unknown and needs to be further investigated.

In summary, our current findings suggest that MFG-E8 plays an inflammatory role in Ang II associated remodeling at the molecular, cellular and tissue levels. MFG-E8 signaling is a key molecular event for Ang II induced adverse arterial remodeling by promoting increased inflammation, VSMC proliferation, collagen deposition, and elastic fiber fragmentation. Thus, targeting MFG-E8 is a potential molecular approach for the prevention or treatment of adverse arterial remodeling during aging and hypertension.

## Author contributions

L.N. and M. W. conceived and designed experiments and wrote the manuscript; L.N., L. L., W.Z., R.T., J.Z., and P.G.G. performed experiments; L.N., L.L., C.H.M., E.G.L., and M.W. analyzed the data; L.N., L.L., C.L., J.L.L, and E.G.L. interpreted results of experiments; K.R.M. prepared Figure 6; L.N. and M.W. wrote the manuscript; R.E.M. and K.R.M. edited the manuscript; and M. W. approved final version of manuscript.

