## Supplementary material for "The Inflammatory Role of Milk Fat Globule Epidermal Growth Factor VIII in Angiotensin II Induced Arterial Remodeling": Suppl text ad figures

Leng Ni, MD, PhD [1,2*]; Lijuan Liu, MD*, MS[1]; Wanqu Zhu, PhD[1]; Richard Telljohann, MS[1]; Jing Zhang, BS[1]; Robert E. Monticone, MS[1]; Kimberly R. McGraw, BS[1] ; Changwei Liu, MD[2]; Christopher H. Morrell, PhD[1]; Pablo Garrido-Gil, PhD[3,4]; Jose Luis Labandeira-Garcia, MD, PhD [3,4]; Edward G. Lakatta, MD[1]; and Mingyi Wang, MD, PhD[1] *1. Laboratory of Cardiovascular Science, National Institution on Aging, National Institutes of Health, Biomedical Research Center (BRC), 251 Bayview Boulevard, Baltimore, MD 21224, USA. 2. Department of Vascular Surgery, Peking Union Medical College Hospital, Peking Union Medical College & Chinese Academy of Medical Sciences, China. 3. Research Center for Molecular Medicine and Chronic Diseases (CIMUS), IDIS, University of Santiago de Compostela, Spain. 4. Networking Research Center on Neurodegenerative Diseases (CIBERNED), Madrid, Spain.*

**Running title:** Arterial Inflammatory Remodeling

**Number of Pages: 13**

**Total Figures: 6**

*Equal contribution

Correspondence to Mingyi Wang, MD, PhD, FAHA

Laboratory of Cardiovascular Science

National Institution on Aging, National Institutes of Health

Biomedical Research Center (BRC), 251 Bayview Boulevard, Baltimore, MD 21224, USA

Tel: (+1) 410-454-8579

### Materials and Methods

#### Experimental Animals

All experiments were conducted according to the protocols (445-LCS-2022) approved by the National Institute on Aging (NIA) in accordance with the National Institute of Health (NIH) Animal Care and Use Committee. MFG-E8 knock out (KO) mice that were generated, characterized, and genotyped as described previously, were obtained from Dr. Mark Udey at National Cancer Institute (NCI).[1, 2] Transgenic Rip1-Tag2 mice were obtained from the National Cancer Institute (NCI). MFG-E8 KO mice were generated by replacing exons 2 to 6 of the gene encoding MFG-E8 in 129SvJ embryonic cells (ES) cells with a neomycin resistant cassette. Mice were genotyped by PCR using the following primers: Rip1-Tag2, GGACAAACCACAACTAGAATGCAGTG (forward) and CAGAGCAGAATTGTGGAGTGG (reverse); Neo, GCCAGAGGCCACTTGTGTAG; and MFG-E8, CTCTCAGATTCACCTGCTCGTG and CACCGTTCAGGCACAGGCTG. 30-week-old male MFGE8 KO mice (homozygous for lack of MFG-E8) and age-matched WT mice were used in this Ang II infusion study. In addition, 4-, 8-, 20, 50-, and 96-week-old male WT and age-matched KO archival aortic sections or frozen tissue (in graphs, individual data represents the number of the aortic samples) were employed for the supplemental study of age-related arterial remodeling. 10-week-old WT and age-matched KO were utilized for measuring circulating lipid profiles.

10-week-old male, WT, AT1, and AT2 receptor knock out mice (WT, AT1 and AT2 KO, respectively) were provided by Professor Jose Luis Labandeira-Garcia.[3] In brief, 10-month-old male C57Bl/6J mice (WT) were used in the present supplemental study. Mice carrying genetic deletions of the Agtr1 (AT1 KO) gene were obtained from The Jackson Laboratory (Bar Harbor, ME, USA) and Agt2 mice (AT2-KO were generously donated by Dr. Daniel Henrion (University of Angers, Angers, France). [4, 5]

**Morphometric Analysis of Aortic Walls in Untreated Aging WT and KO Mice**

We measured intimal medial thickening (IMT) with aging, by staining archival aortic paraffin sections with hematoxylin and eosin (H&E) from untreated 20-week-old young (WT: n=44; KO: n=40) and 96-week-old aged mice (WT: n=39; KO: n=40). The local renin angiotensin system, including Ang II, AT1 and AT2 receptors, and NF-κB activation, were measured in 8-week-old young (WT: n=4; KO: n=5) and 50-week-old aged mice (WT: n=4; MFG-E8 KO: n=4) using immunohistostaining and Western blotting analyses.

**Analysis of Circulating Total Cholesterol, Triglycerides, and Glucose in Untreated WT and KO Mice**

Six male 10-weeks-old MFG-E8 KO (KO) and 6 age-matched WT mice were fasted for 16 hours before sacrifice. The mice were anaesthetized via intraperitoneal injection with ketamine (100 mg/kg of body weight) and xylazine (5 mg/kg per body weight). The thoracic cavity was accessed by cutting the sternum to expose the heart, after a 22 G needle attached to a 3 mL syringe was used to puncture the apex of the heart and 800-1,000 μL of blood was collected, representing most of the animal's total blood volume, and was expected to be arterial blood. Blood was collected in heparin tubes (BD, Franklin Lakes NJ, Item# 36664) and placed on ice for 15 minutes. Samples were then spun down at 10,000 rpm for 10 minutes and plasma was harvested and stored at −80°C until further use. Total cholesterol, triglycerides and glucose levels were measured from plasma samples according to manufacturer instructions. Assay kits for cholesterol (item# 234-60, lot#

54416), triglycerides (item# 236-660, Lot# 54604) and glucose (item# 235-60, lot# 53909) were obtained from Sekure diagnostics (One Wall Street Burlington, MA 01803 USA), and measured using the Cobas Fara II (Roche Diagnostics Roche Diagnostics Corporation 9115 Hague Rd Indianapolis, IN 46256 USA) chemical analyzer.

### Results

#### Ang II, AT1/AT2, and MFG-E8 Expression during Age-associated Inflammatory Remodeling in Mice

Prior studies indicate that arterial Ang II protein and MFG-E8 are increased in aging rats, nonhuman primates, and human samples.[6, 7] Our current findings demonstrate that aortic MFG-E8 protein was markedly elevated in untreated 50-week-old vs 8-week-old WT but was not detected in KO (**Supplemental Figure IA**). Aortic IMT was markedly increased in untreated WT in 96- vesus 20-week-old, but MFG-E8 deficiency alleviates this age effect (**Supplemental Figure IB**). Notably, aortic Ang II protein was significantly increased in untreated aging WT mice, but MFG-E8 deficiency significantly reduced this effect (**Supplemental Figure IC**).

In addition, immunostaining demonstrated that the Ang II receptor AT1 was markedly increased in the aortic wall of untreated WT 96- versus 4-week-old, but the absence of MFG-E8 significantly abolished this age effect; and the Ang II receptor AT2 was markedly decreased in WT aging aortic walls, but the absence of MFG-E8 significantly mitigated these effects (**Supplemental Figure IIA**). Western blot analysis further confirmed the aging effect on AT1 and AT2 expression (50- versus 8-week-old) (**Supplemental Figure IIB**). Interestingly, immunostaining and Western blotting analysis demonstrated that aging markedly increased the master inflammatory transcription factor p-NF-κB p65 in WT, but MFG-E8 deficiency significantly mitigated this age effect (**Supplemental Figure III**).

Importantly, Western blotting analysis demonstrated that the aortic Ang II receptor AT1 was markedly increased in WT after Ang II infusion, but the absence of MFG-E8 significantly reduced this effect (**Supplemental Figure IVA**). However, the Ang II receptor, AT2, was markedly decreased in the aortic walls of both WT and MFG-E8 KO after Ang II infusion. (**Supplemental Figure IVA**). Western blotting analysis of aortic protein from 10-week-old male AT1 and AT2 receptor knock out mice indicated that aortic MFG-E8 protein levels were significantly decreased in AT1 KO, but these levels were not altered in AT2 KO mice (**Supplemental Figure IVB**).

#### Relative Changes in SBP in Ang II Infused Mice

Analysis of the dynamic percentage change in SBP showed there was not a significantly difference in Ang II infused KO versus WT animals (**Supplemental Figure V**).

#### Circulating Cholesterol, Triglyceride, and Glucose Profile in Mice

To determine the profile of total cholesterol, triglyceride, and glucose, sera from 10-week-old KO and 10-week-old WT mice were utilized. Results showed that both fasting cholesterol and triglyceride levels were significantly decreased in KO vs WT, but there was no significant difference in fasting glucose between WT and KO animals (**Supplemental Figure VI**).

**Supplemental Figure Legends**

**Supplemental Figure I. MFG-E8 signaling linked to the age-associated Ang II expression and aortic intimal medial thickening in mice. A.** Representative Western blots of aortic MFG-E8 (left panel). Quantitative data (right panel) of Western blots shows aortic MFG-E8 protein abundance ($p<0.01$ for main age effect, $p<0.0001$ for main genotype effect, $p<0.01$ for age x genotype, by two-way ANOVA). Graph shows mean ± SEM combined with individual data points for KO and WT mice. ***=$p<0.001$ by Bonferroni post-hoc tests following two-way ANOVA with the factors of age and genotype **B.** Morphometric analysis of the aortic intimal medial thickness (IMT) shows IMT ($p<0.0001$ for main age effect, by two-way ANOVA). Graph shows mean ± SEM combined with individual data points for KO and WT mice. **=$p<0.01$ by Bonferroni post-hoc tests following two-way ANOVA. **C.** Representative photomicrographs of immunostaining of aortic Ang II. Morphometric analysis of Ang II immunostaining in the aortic wall shows Ang II abundance ($p<0.0001$ for main age effect, $p<0.0001$ for main genotype effect, $p<0.0001$ for age x genotype, by two-way ANOVA). Graph shows mean ± SEM combined with individual data points for KO and WT mice. ***=$p<0.001$; ****=$p<0.0001$ by Bonferroni post-hoc tests following two-way ANOVA with the factors of treatment and genotype in mice. L=lumen; M=media. Scale bar= 100μm.

**Supplemental Figure II. The age-associated changes of aortic Ang II receptors, AT1 and AT2, expression in mice. A.** Representative photomicrographs of immunostaining of aortic the Ang II AT1 receptor (upper panels). Morphometric analysis of the AT1 immunostaining in the aortic wall shows AT1 abundance ($p<0.01$ for main age effect, $p<0.001$ for main genotype effect, by two-way ANOVA). Graph show mean ± SEM combined with individual data points for KO and WT mice.

*=p<0.05; ***=p<0.001; ****=p<0.0001 by Bonferroni post-hoc tests following two-way ANOVA with the factors of treatment and genotype in mice. Representative photomicrographs of immunostaining of the aortic Ang II AT2 receptor (lower panels). Morphometric analysis shows AT2 abundance (p<0.0001 for main age effect, by two-way ANOVA). Graph shows mean ± SEM combined with individual data points for KO and WT mice. *=p<0.05; ***=p<0.001; ****=p<0.0001 by Bonferroni post-hoc tests following two-way ANOVA with the factors of treatment and genotype in mice. **B.** Representative Western blots of AT1 and AT2 (right panels). Western blotting analysis shows AT1 abundance (middle panel) (p<0.0001 for main age effect, p<0.0001 for main genotype effect, p<0.0001 for age x genotype, by two-way ANOVA). Graph show mean ± SEM combined with individual data points for KO and WT mice. **=p<0.01; ****=p<0.0001 by Bonferroni post-hoc tests following two-way ANOVA with the factors of treatment and genotype in mice. Western blotting analysis shows AT2 abundance (right panel) (p<0.05 for main age effect, p<0.01 for main genotype effect, by two-way ANOVA). Graph shows mean ± SEM combined with individual data points for KO and WT mice. *=p<0.05; *=p<0.01 by Bonferroni post-hoc tests following two-way ANOVA with the factors of treatment and genotype in mice. L=lumen; M=media. Scale bar= 100µm.

**Supplemental Figure III. The age-associated activation of NF-κB in the aortic wall of mice. A.** Representative photomicrographs of immunostaining of aortic p-NF-κB p65. Morphometric analysis of the percentage of p-NF-κB p65 stained nuclei in the aortic wall shows NF-κB activation (p<0.001 for main age effect, p<0.0001 for main genotype effect, p<0.001 for age x genotype, by two-way ANOVA). Graph shows mean ± SEM combined with individual data points for KO and WT mice. ****=p<0.0001 by Bonferroni post-hoc tests following two-way ANOVA with the factors of treatment and genotype in mice. **B.** Representative Western blots of NF-κB p65.

Western blotting analysis shows p-NF-κB p65 abundance (right panel) ($p<0.0001$ for main age effect, $p<0.0001$ for main genotype effect, $p<0.0001$ for age x genotype, by two-way ANOVA). Graph shows mean ± SEM combined with individual data points for KO and WT mice. *=$p<0.05$; ****=$p<0.0001$ by Bonferroni post-hoc tests following two-way ANOVA with the factors of treatment and genotype in mice. L=lumen; M=media. Scale bar= 100μm.

**Supplemental Figure IV. Effects of Ang II receptors on MFG-E8 expression in the aortic wall of mice. A.** Representative Western blots of AT1 receptors (left panel). Western blotting analysis shows AT1 abundance (middle panel) ($p<0.0001$ for main Ang II treatment effect, $p<0.0001$ for main genotype effect, $p<0.0001$ for treatment x genotype, by two-way ANOVA). Graph (right panel) show mean ± SEM combined with individual data points for KO and WT mice. ***=$p<0.001$; ****=$p<0.0001$ by Bonferroni post-hoc tests following two-way ANOVA with the factors of treatment and genotype in mice. **B.** Representative Western blots of AT2 receptor (right panel). Western blotting analysis shows AT2 abundance (middle panel) ($p<0.001$ for main Ang II infusion effect, $p<0.05$ for main genotype effect, by two-way ANOVA). Graph (right panel) shows mean ± SEM combined with individual data points for KO and WT mice. **=$p<0.01$ by Bonferroni post-hoc tests following two-way ANOVA with the factors of treatment and genotype in mice. **C**. Representative Western blots of aortic MFG-E8 from 10-week-old AT1 and AT2 KO mice. Western blotting analysis shows MFG-E8 abundance ($p<0.001$ for the genotype effect, by one-way ANOVA). Graph shows mean ± SEM combined with individual data points for KO and WT mice. ****=$p<0.0001$ by Bonferroni post-hoc tests following one-way ANOVA in mice.

**Supplemental Figure V. Dynamic percentage changes in SBP in mice.** Dynamic percentage changes in mouse systolic blood pressure (SBP) as measured by tail-cuff method; plot shows mean

± SEM for KO and WT mice (n=5/group) with Ang II or saline infusion over time, $p>0.05$, analyzed by repeated two-way ANOVA)

**Supplemental Figure VI. Circulating cholesterol, triglyceride, and glucose profile in mice.** Plasma cholesterol (left panel), triglyceride (middle panel), and glucose (right panel). Graph shows mean ± SEM combined with individual data points for KO and WT mice. Two samples unpaired T-TEST: ** =$p<0.01$; ****=$p<0.0001$.

7. Fu Z, Wang M, Gucek M, Zhang J, Wu J, Jiang L, Monticone RE, Khazan B, Telljohann R, Mattison J, Sheng S, Cole RN, Spinetti G, Pintus G, Liu L, Kolodgie FD, Virmani R, Spurgeon H, Ingram DK, Everett AD, Lakatta EG, Van Eyk JE. Milk fat globule protein

epidermal growth factor-8: A pivotal relay element within the angiotensin ii and monocyte chemoattractant protein-1 signaling cascade mediating vascular smooth muscle cells invasion. *Circulation research*. 2009;104:1337-1346

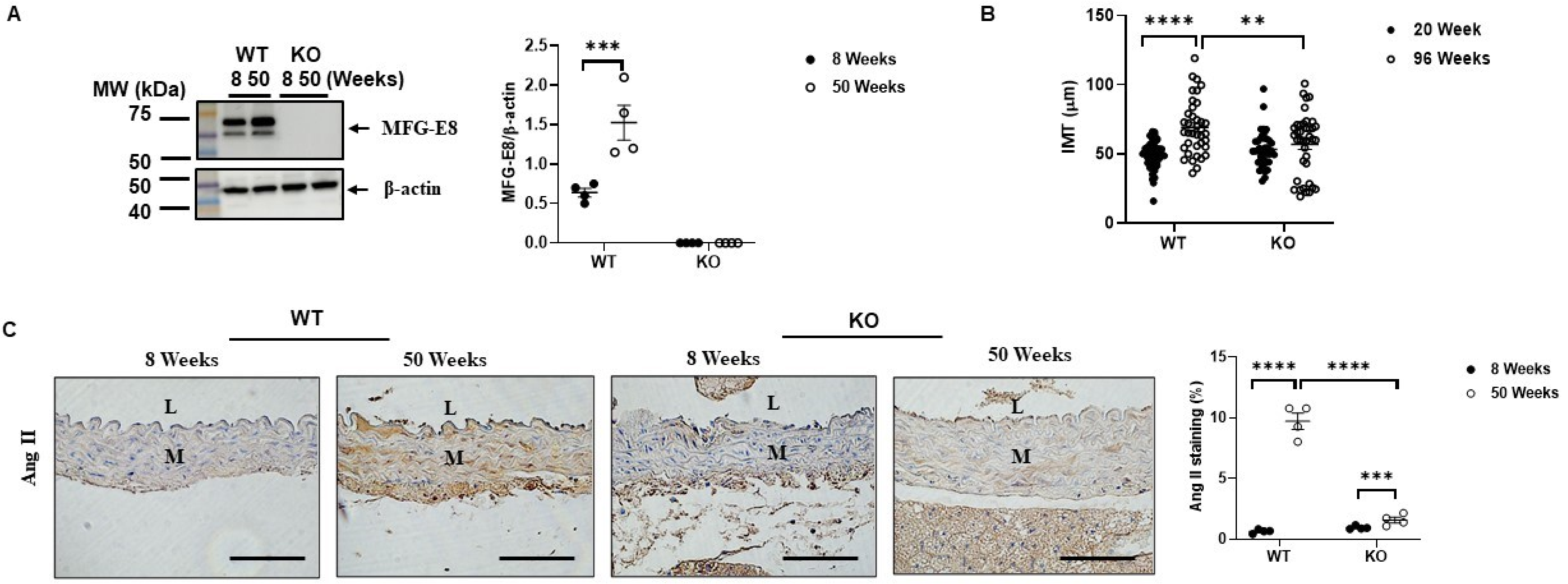


Suppl Figure I

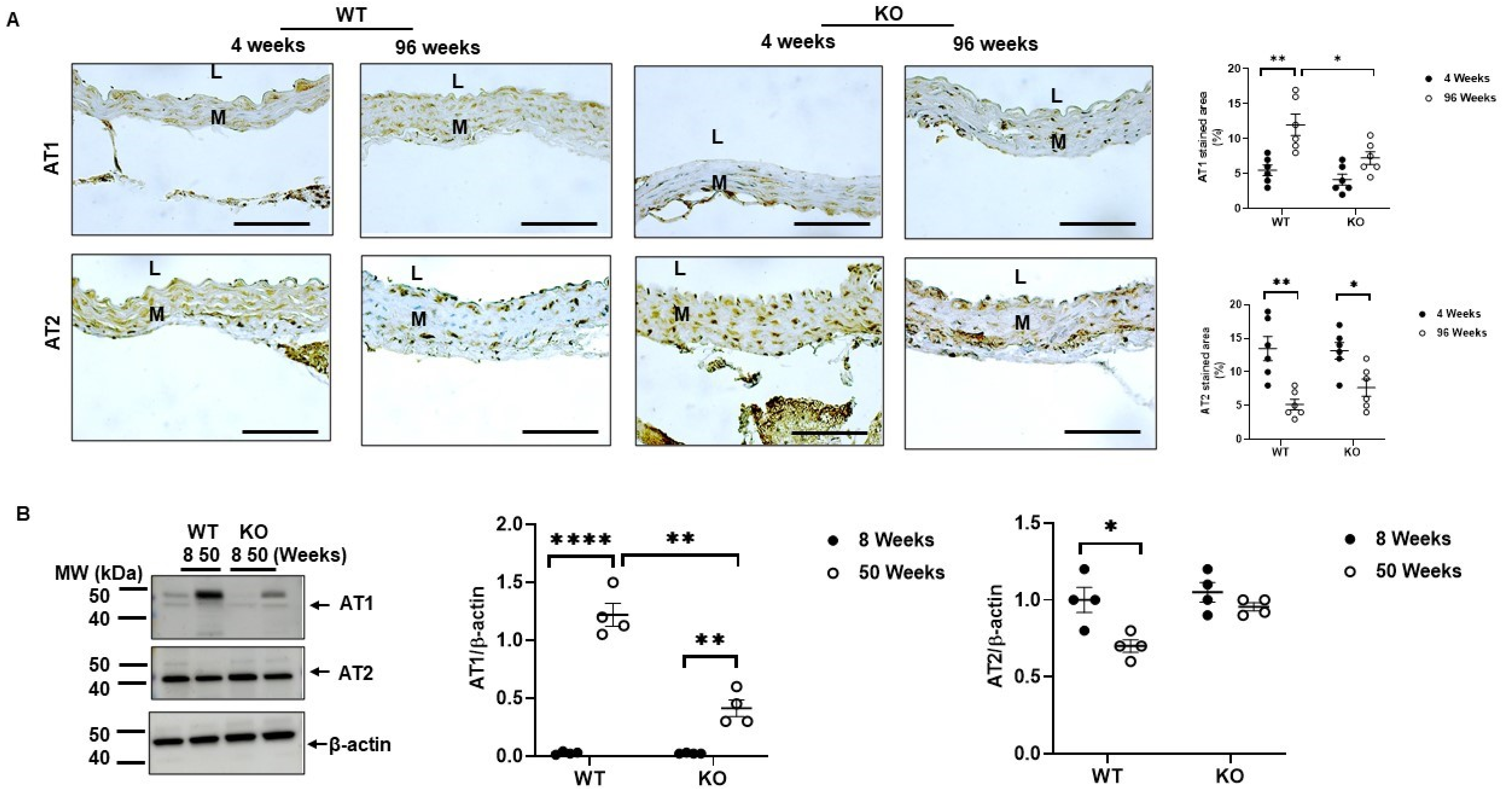


Suppl Figure II

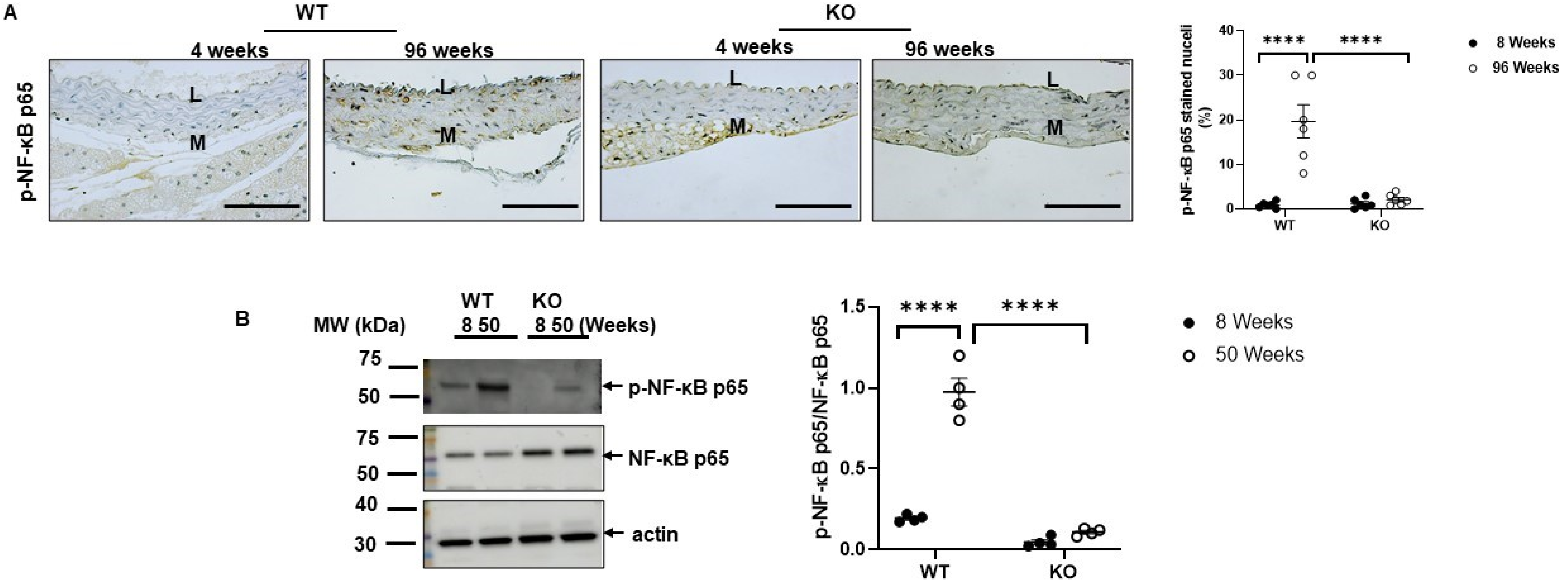


Suppl Figure III

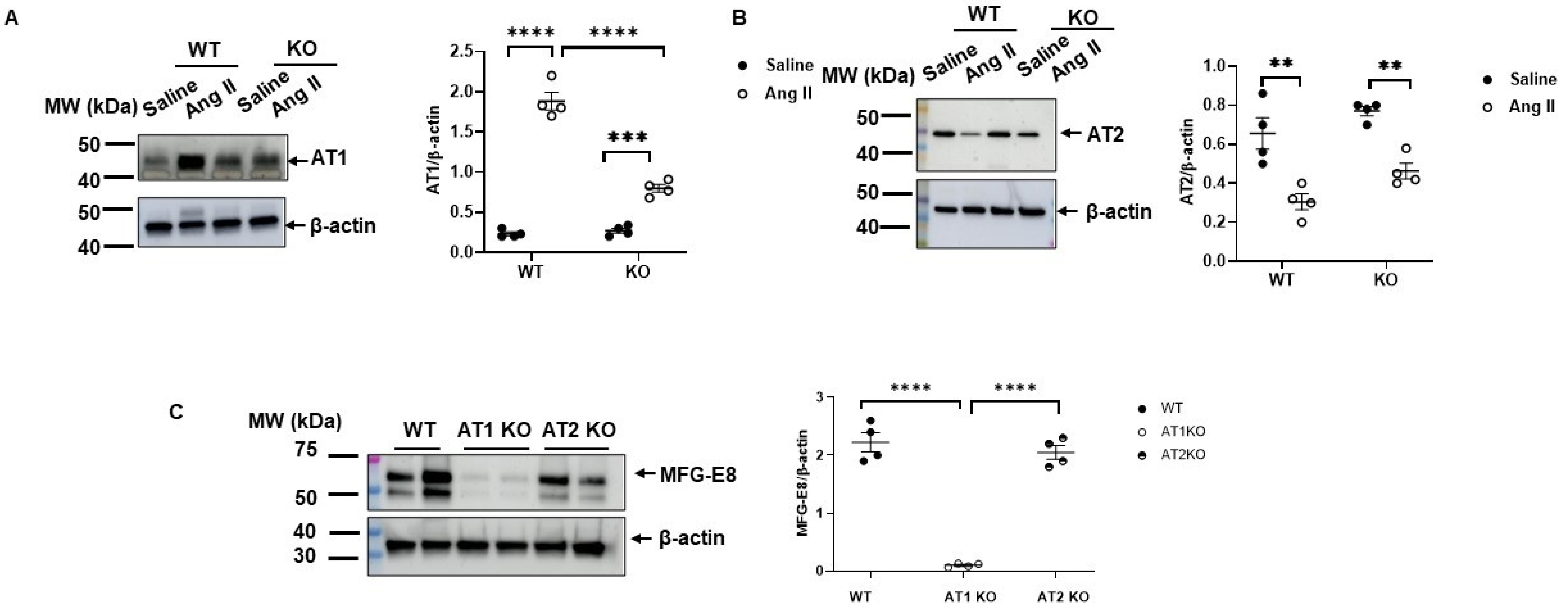


Suppl. Figure IV

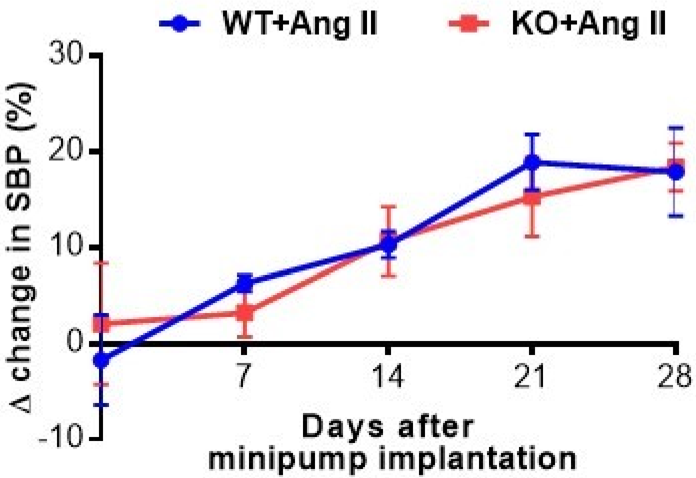


Suppl. Figure V

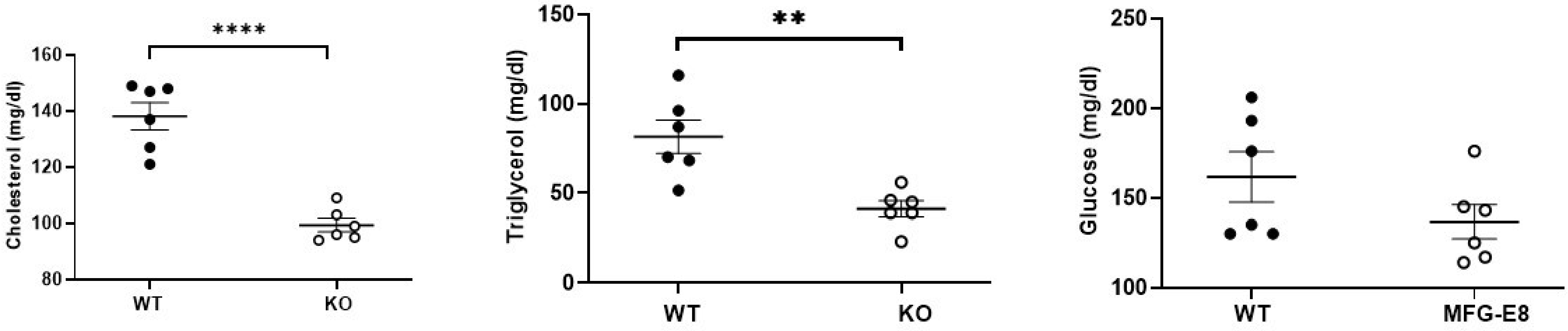


Suppl. Figure VI
